# Fungal pathogens exposed – genomic and phenotypic insights into *Candida auris* and its relatives of the *Candida haemulonii* species complex

**DOI:** 10.1101/2024.05.16.594571

**Authors:** Auke W. de Jong, Sander Boden, Annemarie Zandijk, Alexandra M. Kortsinoglou, Bert Gerrits van den Ende, Elaine C. Francisco, Podimata Konstantina Nefeli, Vassili N. Kouvelis, Miaomiao Zhou, Ferry Hagen

## Abstract

*Candida auris* showed the world the ability of fungi to rapidly emerge as an urgent threat to public health. Close relatives of the *Candida haemulonii* complex exhibit also a similar multi-drug resistant nature and are increasingly reported as nosocomial pathogens. Here, we analyze both complete genomes assemblies and extensive phenotypic data for the five *C. auris* clades and pathogenic and non-pathogenic species related to the *C. haemulonii* species complex. First, we resolved the phylogeny of the species complex. Next, comparing *C. auris* to its pathogenic and non-pathogenic relatives we identified a pathogenic potential for the whole *C. haemulonii* species complex by shared gene content and phenotypic traits linked to drug resistance and virulence. *In vivo* virulence assays using the *Galleria mellonella* fungal infection model show that *C. auris* strains are significantly more virulent than any of the sibling species in the *C. haemulonii* complex. Phenotypic analysis links the increased virulence of *C. auris* to a more stress resistant phenotype compared to its siblings.

## Introduction

Fungi are on the rise. Climate change, globalization, habitat disturbance, and an increasing population of immunocompromised patients all contribute to the spread and rising incidence of fungal infections (Fisher et al. 2018; Geddes-McAlister et al., 2019; Garcia-Solache et al., 2010). Each year, over 300 million people worldwide are affected by serious fungal infections, resulting annually in 2.5 million deaths (Denning, 2024). These numbers are on par with the number of deaths caused by well-known bacterial and parasite pathogens, such as tuberculosis and malaria (Brown et al., 2012). The economic burden of fungal diseases in the US alone was estimated at $11.5 billion in 2019 (Benedict et al., 2022). This is likely an underestimation due to persisting underdiagnoses and underreporting. Nevertheless, fungi are notoriously neglected as a serious threat to global health making them an unrecognized pandemic (Nat Microbiol, 2017). In 2022, the World Health Organization (WHO) released the fungal priority pathogens list (WHO FPPL), a significant milestone in the recognition of fungal infections (WHO, 2022).

*Candida* yeasts are the primary cause of hospital-acquired fungal infections, estimated to affect >1.5 million people each year (Denning, 2024; Pfaller et al., 2019; Bassetti et al., 2019). Therefore, *Candida* contributes considerably to morbidity, mortality, and economic losses caused by fungal infections. While *Candida albicans* is still recognized as the main cause of candidiasis, other *Candida* species are on the rise (Pfaller et al., 2019; Stavrou et al., 2019). Most of these emerging species show reduced sensitivity to one or more of the three antifungal drugs classes used in clinical settings. Increased resistance combined with the misidentification of emerging species by commercial biochemical systems and standard laboratory methods makes adequate treatment increasingly challenging (Kathuria et al., 2015; Castanheira et al., 2013; Gow et al., 2022).

*Candida auris* rapidly became the most notorious emerging fungal pathogen. This multi-drug resistant yeast has conquered nosocomial environments all over the world by storm. The US Centers for Disease Control and Prevention even classified *C. auris* as one of five microbes that are the most urgent threat to public health, and the WHO recently listed *C. auris* as a ‘critical priority fungal pathogen’ (WHO, 2022; CDC, 2019). *C. auris* has the unique ability to persistently colonize the hospital environment and the host skin resulting in high transmission rates and subsequent outbreaks (de Jong & Hagen, 2019). Nevertheless, the unprecedented global spread of *C. auris* remains a mystery. Genomic analyses revealed the near simultaneous, but independent, emergence of four distinct clades on different continents (South Asian (I), East Asian (II), African (III), and South American (IV)) (Lockhart et al., 2017; Chow et al., 2020). A minor fifth and more recent sixth clade were also described (Chow et al., 2018; Suphavilai et al., 2024).

Despite *C. auris* being the most worrisome emerging fungal pathogen, more yeasts are on the rise. Phylogenetic studies revealed that *C. auris* is closely related to the *Candida haemulonii* complex (Muñoz et al., 2018; Gade et al., 2020; Francisco et al., 2023). This complex contains other emerging nosocomial pathogens such as *Candida haemulonii*, *Candida duobushaemulonii*, and *Candida pseudohaemulonii* (Muñoz et al., 2018; Cendejas-Bueno et al., 2012). In recent years, additional members have been identified including new pathogens such as *Candida vulturna* and *Candida khanbhai*, but also the non-pathogenic species *Candida chanthaburiensis*, *Candida heveicola*, *Candida konsanensis*, *Candida metrosideri*, *Candida ohialehuae*, and *Candida ruelliae* (Jackson et al., 2019; Klaps et al., 2020; Sipiczki & Tap, 2016; de Jong et al., 2023). This independent evolution of pathogenicity within the same complex is observed throughout the genus *Candida* (Stavrou et al., 2019; Gabaldón et al., 2016; Rokas, 2022). The mix of closely related pathogens and non-pathogens within the *C. haemulonii* complex suggests that members within this complex harbor traits that pre-adapt them to human pathogenicity (Rokas, 2022). Nevertheless, pathogens of the *C. haemulonii* complex differ substantially in their virulence (Fakhim et al., 2018; Muñoz et al., 2020). This observation is similar in other *Candida* complexes. For example, *C. albicans* is much more virulent than its sibling species *C. dubliniensis* (Singh-Babak et al., 2021), and the common pathogen *Nakaseomyces glabratus* is closely related to the non-pathogenic species *Nakaseomyces castellii* (Gabaldón et al., 2016).

Notably, variation in pathogenicity is not restricted between complexes and species but is even present among strains of the same species (Rokas, 2022). *C. auris* abundantly demonstrates this strain heterogeneity. Particularly, clade-specific differences in virulence and antifungal resistance are found. For example, Clade II and Clade V strains typically cause relatively harmless ear infections, while strains from other clades cause life-threatening invasive blood infections (Chow et al., 2018; Welsh et al., 2019). Strain specific virulence differences of *C. auris* have also been observed in both *Galleria mellonella* and mice *in vivo* infection models (Fakhim et al., 2018; Muñoz et al., 2020; Forgács et al., 2020). In addition, resistance levels vary significantly between *C. auris* strains, with some being pan-resistant and others totally susceptible (Chow et al., 2020; de Jong et al., 2022). Most clinical strains are resistant against fluconazole, but resistance to amphotericin B is common too (Chow et al., 2020). Species of the *C. haemulonii* complex have a similar resistance pattern displaying intrinsic resistance against azoles and amphotericin B (Cendejas-Bueno et al., 2012). While resistance mechanisms in *C. auris* are starting to be understood, those used by other *C. haemulonii* complex members still need to be elucidated (Carolus et al., 2021; Rybak et al., 2022). Finally, initial studies observed strain-specific variability for *C. auris* in other virulence factors such as lytic enzyme production, stress resistance and biofilm production (de Jong & Hagen, 2019). However, data on phenotypic differences between *C. auris* clades is highly fragmented and little to none is known about differences with other species of the *C. haemulonii* complex.

Although the mechanisms of infection of the well-known *Candida* pathogens are starting to be unveiled, it remains unclear what exactly makes a virulent phenotype. It is to be expected that infection-relevant traits are more pronounced in human pathogens compared to their closest related non-pathogenic relatives. While comparative genomic efforts provided an initial idea of the mechanisms behind the pathogenic success of *C. auris* and some of its relatives, only few strains were used and a complete representation of the *C. haemulonii* complex is lacking (Muñoz et al., 2018). Here we combine genomic and extensive phenotypic data for both pathogenic and non-pathogenic members of the *C. haemulonii* complex to elucidate phylogenetic relationships and gain novel insights into the evolution of pathogenesis within this increasingly important species complex.

## Methods

### Strains and culture

Details of the strains used in this study are listed in Table 1. *C. auris* strains were selected as follows: two strains per clade, representing the four clades that are specific to each geographic region, and one strain representing the rare fifth clade. Strains of other members of the *C. haemulonii* complex and close relatives were selected to represent both clinical and environmental strains of each species when possible (Table 1). Strains were maintained at - 80°C in yeast extract 1%, peptone 2%, dextrose 2% (YPD) medium with 25% glycerol. Before use strains were subcultured onto YPD with 1.5% agar at 25°C for 24–48h. A single colony was picked and put onto YPD slants that were kept at room temperature as a stock culture for inoculation of future experiments. Specific culture conditions for each experiment are described below.

**Table 1.**
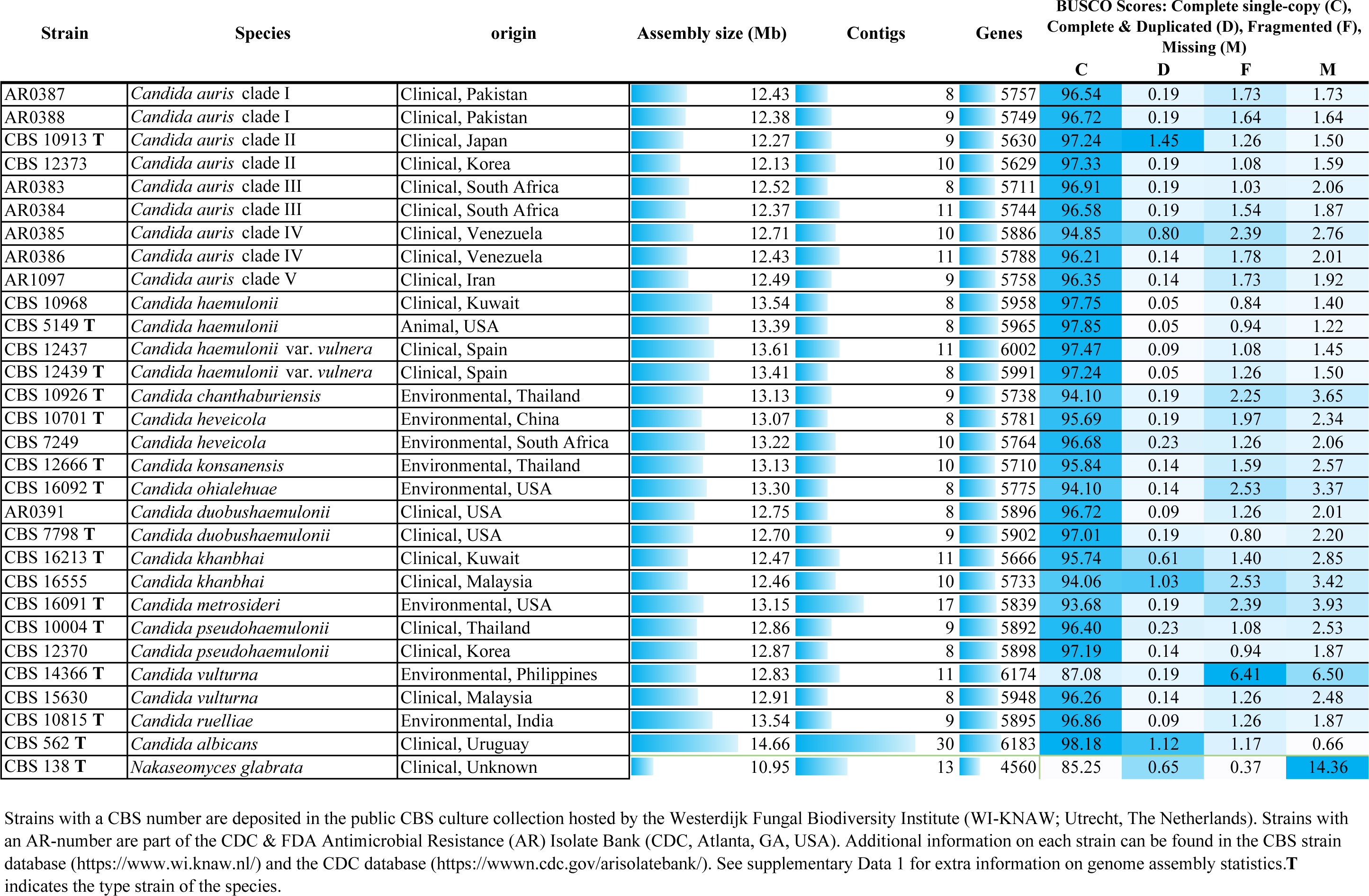
*Candida auris* and *C. haemulonii* species complex genome assembly statistics. Strains with a CBS number are deposited in the public CBS culture collection hosted by the Westerdijk Fungal Biodiversity Institute (WI-KNAW; Utrecht, The Netherlands). Strains with an AR-number are part of the CDC & FDA Antimicrobial Resistance (AR) Isolate Bank (CDC, Atlanta, GA, USA). Additional information on each strain can be found in the CBS strain database (https://www.wi.knaw.nl/) and the CDC database (https://www.n.cdc.gov/arisolatebank/). All assemblies reported in this study are deposited in the NCBI GenBank Database; Bioproject number: PRJNA1002224, PRJNA1003053 and PRJNA1050609. See supplementary Data 1 for extra information on genome assembly statistics. **T** indicates the type-strain of the species.

| Strain | Species | origin | Assembly size (Mb) | Contigs | Genes | BUSCO Scores: Complete single-copy (C), Complete & Duplicated (D), Fragmented (F), Missing (M) |  |  |  |
| --- | --- | --- | --- | --- | --- | --- | --- | --- | --- |
|  |  |  |  |  |  | C | D | F | M |
| AR0387 | *Candida auris* clade I | Clinical, Pakistan | 12.43 | 8 | 5757 | 96.54 | 0.19 | 1.73 | 1.73 |
| AR0388 | *Candida auris* clade I | Clinical, Pakistan | 12.38 | 9 | 5749 | 96.72 | 0.19 | 1.64 | 1.64 |
| CBS 10913 T | *Candida auris* clade II | Clinical, Japan | 12.27 | 9 | 5630 | 97.24 | 1.45 | 1.26 | 1.50 |
| CBS 12373 | *Candida auris* clade II | Clinical, Korea | 12.13 | 10 | 5629 | 97.33 | 0.19 | 1.08 | 1.59 |
| AR0383 | *Candida auris* clade III | Clinical, South Africa | 12.52 | 8 | 5711 | 96.91 | 0.19 | 1.03 | 2.06 |
| AR0384 | *Candida auris* clade III | Clinical, South Africa | 12.37 | 11 | 5744 | 96.58 | 0.19 | 1.54 | 1.87 |
| AR0385 | *Candida auris* clade IV | Clinical, Venezuela | 12.71 | 10 | 5886 | 94.85 | 0.80 | 2.39 | 2.76 |
| AR0386 | *Candida auris* clade IV | Clinical, Venezuela | 12.43 | 11 | 5788 | 96.21 | 0.14 | 1.78 | 2.01 |
| AR1097 | *Candida auris* clade V | Clinical, Iran | 12.49 | 9 | 5758 | 96.35 | 0.14 | 1.73 | 1.92 |
| CBS 10968 | *Candida haemulonii* | Clinical, Kuwait | 13.54 | 8 | 5958 | 97.75 | 0.05 | 0.84 | 1.40 |
| CBS 5149 T | *Candida haemulonii* | Animal, USA | 13.39 | 8 | 5965 | 97.85 | 0.05 | 0.94 | 1.22 |
| CBS 12437 | *Candida haemulonii* var. *vulnera* | Clinical, Spain | 13.61 | 11 | 6002 | 97.47 | 0.09 | 1.08 | 1.45 |
| CBS 12439 T | *Candida haemulonii* var. *vulnera* | Clinical, Spain | 13.41 | 8 | 5991 | 97.24 | 0.05 | 1.26 | 1.50 |
| CBS 10926 T | *Candida chanthaburiensis* | Environmental, Thailand | 13.13 | 9 | 5738 | 94.10 | 0.19 | 2.25 | 3.65 |
| CBS 10701 T | *Candida heveicola* | Environmental, China | 13.07 | 8 | 5781 | 95.69 | 0.19 | 1.97 | 2.34 |
| CBS 7249 | *Candida heveicola* | Environmental, South Africa | 13.22 | 10 | 5764 | 96.68 | 0.23 | 1.26 | 2.06 |
| CBS 12666 T | *Candida konsanensis* | Environmental, Thailand | 13.13 | 10 | 5710 | 95.84 | 0.14 | 1.59 | 2.57 |
| CBS 16092 T | *Candida ohialehuae* | Environmental, USA | 13.30 | 8 | 5775 | 94.10 | 0.14 | 2.53 | 3.37 |
| AR0391 | *Candida duobushaemulonii* | Clinical, USA | 12.75 | 8 | 5896 | 96.72 | 0.09 | 1.26 | 2.01 |
| CBS 7798 T | *Candida duobushaemulonii* | Clinical, USA | 12.70 | 9 | 5902 | 97.01 | 0.19 | 0.80 | 2.20 |
| CBS 16213 T | *Candida khanbhai* | Clinical, Kuwait | 12.47 | 11 | 5666 | 95.74 | 0.61 | 1.40 | 2.85 |
| CBS 16555 | *Candida khanbhai* | Clinical, Malaysia | 12.46 | 10 | 5733 | 94.06 | 1.03 | 2.53 | 3.42 |
| CBS 16091 T | *Candida metrosideri* | Environmental, USA | 13.15 | 17 | 5839 | 93.68 | 0.19 | 2.39 | 3.93 |
| CBS 10004 T | *Candida pseudohaemulonii* | Clinical, Thailand | 12.86 | 9 | 5892 | 96.40 | 0.23 | 1.08 | 2.53 |
| CBS 12370 | *Candida pseudohaemulonii* | Clinical, Korea | 12.87 | 8 | 5898 | 97.19 | 0.14 | 0.94 | 1.87 |
| CBS 14366 T | *Candida vulturna* | Environmental, Philippines | 12.83 | 11 | 6174 | 87.08 | 0.19 | 6.41 | 6.50 |
| CBS 15630 | *Candida vulturna* | Clinical, Malaysia | 12.91 | 8 | 5948 | 96.26 | 0.14 | 1.26 | 2.48 |
| CBS 10815 T | *Candida ruelliae* | Environmental, India | 13.54 | 9 | 5895 | 96.86 | 0.09 | 1.26 | 1.87 |
| CBS 562 T | *Candida albicans* | Clinical, Uruguay | 14.66 | 30 | 6183 | 98.18 | 1.12 | 1.17 | 0.66 |
| CBS 138 T | *Nakaseomyces glabrata* | Clinical, Unknown | 10.95 | 13 | 4560 | 85.25 | 0.65 | 0.37 | 14.36 |

### Nanopore sequencing, genome assembly and gene annotation

For long-read nanopore sequencing DNA was extracted using an established cetyltrimethylammonium bromide (CTAB) DNA extraction protocol optimized to yield high quantity and quality genomic DNA as previously described (Navarro-Muñoz et al., 2019). DNA concentrations were quantified by Qubit fluorometric method (ThermoFisher, Waltham, MA, USA). Genomic libraries were prepared and barcoded using the ligation sequencing kit (SQK-LSK109; ONT, Oxford, UK) and native barcoding kit (EXP-NBD104; ONT) and run on a MinION flow cell (FLO-MIN106D R9.4; ONT) according to the manufacturer’s instructions. Basecalling of raw data was performed with Guppy v6 (Wick et al., 2019). For raw read data, quality control was done using NanoPlot v1.39.0 (De Coster et al., 2018), before and after read filtering/trimming by NanoFilt v2.8.0 (De Coster et al., 2018). The filtered reads were then assembled using Flye v2.9.2 (Kolmogorov et al., 2019) followed by two rounds of polishing. Each assembly was assessed using Quast v5.0.2 (Gurevich et al., 2013) to generate statistics of assembly quality. Genome assemblies were manually checked, and mitochondrial genomes were corrected for redundancy prior data deposition in the NCBI Genome database (see Data availability statement). Draft assemblies were annotated using Helixer v0.3.1 for *de novo* prediction of gene structure combining deep learning and a hidden Markov Model (Holst et al., 2023). A specific fungi training model which includes *C. auris* and *C. haemulonii* references (fungi_v0.3_a_0100), was set and run using the general recommendations provided by the authors. Annotation quality was assessed by BUSCO v5.3.0 in protein mode (Simão et al., 2015).

### Phylogenomic analysis and comparative genomics

Protein sequences were extracted from the Helixer annotations using the gffread utility v0.12.7. Orthology and phylogenetic analysis were performed by OrthoFinder v2.5.3 (Emmes et al., 2019). In addition, average nucleotide identity was calculated using OrthoANI (Lee et al., 2016). Conservation of resistance and virulence genes was compared using a list of genes associated with various resistance mutations and virulence traits compiled from the literature (Supplementary Data 2). Orthologous clusters containing these genes were extracted from the OrthoFinder data using the respective protein sequence from *C. auris* reference strain B8441 (=AR0387) (https://www.candidagenome.org/; Supplementary Data 2). Sums of gene counts from these extracted clusters were calculated per strain, per trait. The summed gene counts were visualized along with the OrthoFinder phylogeny and phenotypic data using ITOL (Letunic et al., 2021).

### Phenotypical analysis

All strains were phenotypically characterized based on standard physiological procedures as previously described (Kurtzman et al., 2011). Fermentation and assimilation of carbohydrates were performed in liquid media at 25°C up to 21 days. Assimilation of nitrogen compounds was assessed with the auxanographic method. Additionally, specific virulence related characteristics like stress resistance, lytic enzyme production and biofilm formation (de Jong & Hagen., 2019) were determined by the methods described below.

### Stress resistance assays

For the stress resistance assays, strains were grown in YPD for 18h at 30°C in liquid shaken culture at 200rpm, unless stated otherwise. Cells were collected by centrifugation and washed twice using phosphate-buffered saline (PBS; without Ca, Mg; pH 7.3–7.5; Lonza, Basel, Switzerland). Ten-fold serial dilutions were made and 5µl of each dilution was spotted onto YPD agar plates, which contained the following compounds: 1M CaCl_2_, 300 μg/ml Calcofluor white, 300 μg/ml Congo Red, 10% glycerol, 1.75M NaCl (obtained from Merck, Darmstadt, Germany) and 15 mM β-mercapto-ethanol, 15mM caffein, 12 mM hydrogen peroxide (H_2_O_2_), 1M KCl 0.05% SDS (obtained from ThermoFisher) or YPD adjusted to pH3 with 37% HCl solution or pH8 with 5M Tris buffered saline. After 48h at 37°C, growth was compared to the control YPD plates. The growth of each strain was rated qualitatively on a scale of 0 to 5 (see Supplementary Data 4 for assignment of the scale).

Hydrogen peroxide resistance was tested in liquid culture using a 96-wells format in which cells were inoculated at different H_2_O_2_ concentrations (0, 1, 5, 7.5, 10, 15, 25, and 50mM). Cells were enumerated using a Cellometer X2 cell counter (Nexcelom Bioscience, Lawrence, MA, USA) and resuspended in yeast nitrogen base supplemented with 2% glucose (Difco; BD, Franklin Lakes, NJ, USA), to obtain a concentration of 2.0 × 10^6^ cells/ml. The 96-wells plates were filled with 50µl of 2× the indicated concentrations of H_2_O_2_. Next, 50 µl of the cell suspensions was added to obtain the indicated H_2_O_2_ concentrations and a final cell concentration of 1.0 × 10^6^ cells/ml. Strains were inoculated in quadruplicate. Plates were incubated at 25°C and 37°C. After 48h the concentration of H_2_O_2_ at which growth was still observed was determined using a SPECTROstar Nano spectrophotometer (BMG Labtech, Ortenberg, Germany) at 600nm. The experiments were repeated twice.

### Biofilm formation

Biofilm formation was evaluated using standardized methods described previously (Ramage et al., 2001), with slight modifications. Briefly, cell suspensions were made in YNB with 2% glucose to a concentration of 1.0 × 10^6^cells/ml. On two separate occasions, 4 biofilms of each *Candida* species were grown in Cellstar flat-bottomed 96-well plates (Greiner bio-one, Alphen aan den Rijn, The Netherlands). One-hundred µl of cell suspension was added to each well and the plates were incubated at 25°C and 37°C for 24h. After washing the biofilms an XTT (2,3-bis(2-methoxy-4-nitro-5-sulfo-phenyl)-2H-tetrazolium-5-carboxanilide-reduction assay was carried out to measure biofilm activity as a semiquantitative measure of biofilm formation. To this end, the Cell proliferation Kit II (XTT) (Roche, Basel, Switzerland) was used. Fifty µl of the XTT labelling mixture was added to the washed biofilms and incubated in the dark for 1h at 35°C. After incubation the supernatant was transferred to a clean 96-wells plate and biofilms were washed with 50µl PBS to dissolve left-over XTT product. This was added to the first 50µl in the clean plate to end up with 100µl aliquots, which were measured at 492nm using an automated plate reader. The plate with biofilms was then air dried and additionally used to quantify biofilm biomass using a crystal violet assay as described elsewhere (Jose et al., 2010).

### Enzymatic activity

Enzymatic activity was evaluated using different culture media to measure phospholipase (Price et al., 1982), lipase (Buzzini & Martini, 2002), esterase (Slifkin et al., 2000), secreted aspartyl proteinase (SAP)(Crandall & Edwards, 1987), and hemolysin (Luo et al., 2001) activity. To this end, strains were inoculated in YNB broth with 2% glucose for 18h at 30°C, and 200rpm. Cells were collected and washed twice with PBS by centrifugation at 3000 × *g* for 2 min. Next, the samples were diluted in PBS to a final concentration of 1.0 × 10^8^ cells/ml. 5 µl cell suspension was spotted in triplicates onto the assay plates and incubated at 25°C and 37°C for 2 days (hemolysin), 5 days (phospholipase and SAP) or 7 days (lipase). The enzymatic activity index value (Pz) was calculated as (1 - (diameter colony / diameter precipitation zone)). Hemolysin activity was divided into β activity represented by a translucent halo around the colony defined as the hemolytic index (Hi) and α activity represented by a black-greenish ring at the periphery of the distinctive translucent halo defined as the peripheral hemolytic index (Hp)(Wan et al., 2015). All media were prepared according to the previously described protocols except for the blood (hemolysin) and tributyrin (lipase) agar plates, which were prepared with 5% fresh human blood and 0.1% tributyrin, respectively. Each experiment was repeated twice.

### Antifungal susceptibility testing

*In vitro* antifungal susceptibility of each strain was determined using the EUCAST broth microdilution method according to protocol E.DEF.7.3.2 (Arendrup et al., 2020). Included antifungals in this test were: amphotericin B (AMB), anidulafungin (AND), micafungin (MCF), 5-flucytosine (5FC), azoles including fluconazole (FLU), itraconazole (ITR), voriconazole (VOR), posaconazole (POS) and isavuconazole (ISA) (all obtained from Sigma Aldrich, St. Louis, MI, USA). Minimal inhibitory concentration (MIC) values were determined after 24h and 48h. The tentative MIC breakpoints given for *C. auris* by the CDC were used as references (CDC, 2018).

### Galleria mellonella survival assays

Survival assays were performed using the *Galleria mellonella* model for fungal infection following previously described protocols (Fuchs et al., 2010). Briefly, *Candida* strains were grown for 18h in YNB supplemented with 2% glucose at 30°C, 200rpm. Cells were collected by centrifugation and washed twice with PBS. After counting with a Cellometer X2 cell counter (Nexcelom Bioscience), the cell suspensions were adjusted to 1.0 × 10^8^ cells/ml in PBS. Larvae were purchased from Kreca Ento-Feed (Ermelo, The Netherlands) and sorted to obtain homogenous groups of 15 larvae without grey markings, weighing approximately 250mg each (de Jong et al., 2022). Larvae were maintained at 25°C in the dark and used within 5 days of receipt. Individual larvae were cleaned with 70% ethanol and inoculated in the left rear proleg with 1.0 × 10^6^ yeast cells (10 µl final inoculum volume) using a 10-µl Hamilton syringe fitted with a 26-gauge point style 2 needle. Two control groups were included: untreated larvae and larvae injected with PBS. Larvae were put in petri dishes without food and incubated at 37°C over a period of 10 days. Viability was scored daily, and dead larvae were removed together with faeces and webbing. The experiment was repeated three times on different occasions. The combined (additive) data from all experiments (n = 45) was used to generate Kaplan–Meier survival plots and analysed using the Mantel-Cox pairwise Log-rank test: P<0.001; ns, P>0.05 (GraphPad Prism v10.0.0, GraphPad Software, Boston, MA, USA).

### Data availability

Yeast strains used in this study have been deposited in the CBS culture collection (hosted at the Westerdijk Fungal Biodiversity Institute, Utrecht, The Netherlands), or are available via the CDC Isolate Bank (hosted at the Centers for Disease Control & Prevention, Atlanta, GA, USA). All genome assemblies have been deposited in the NCBI GenBank repository under the following BioProject numbers: PRJNA1002224, PRJNA1003053 and PRJNA1050609. The individual strains and sequence data accession numbers are provided in Supplementary Data 1.

## Results

### Genome features of the *Candida haemulonii* complex

To determine potential genomic characteristics related to virulence within *C. auris* and the *C. haemulonii* complex, we generated complete genomes of each species (Table 1). Strains of all five *C. auris* clades, and its pathogenic relatives *C. haemulonii*, *C. pseudohaemulonii*, *C. duobushaemulonii*, *C. vulturna* (Navarro-Muñoz et al., 2019), and the recently described clinical species *C. khanbhai* were included (de Jong et al., 2023). Additionally, non-pathogenic relatives including *C. chanthaburiensis*, *C. heveicola*, *C. konsanensis*, *C. metrosideri*, *C. ohialehuae*, and *C. ruelliae* were also sequenced to enable comparative genomic analysis of pathogens and non-pathogens (Table 1). Where possible, two strains of each species were selected to also enable intraspecies genome comparison (Table 1). All genome assemblies were sequenced using nanopore long-read technology. As an outgroup genome assemblies of *C. albicans* CBS 562 and *N. glabratus* CBS 138 were included for comparison.

*C. auris* genomes were organized in 8 to 11 contigs. Like previous highly contiguous genome assemblies generated by long reads, most of the sequenced bases are grouped into 7 contigs and an 8^th^ circular contig (Muñoz et al., 2018). This is in line with the presence of 7 chromosomes and a circular mitochondrial genome reported by previous studies for *C. auris* (Muñoz et al., 2018; Muñoz et al., 2021; Misas et al., 2020).

For most other members of the *C. haemulonii* species complex, only Illumina or no genome data at all was available. Here we generated highly contiguous genomes for these species with the amount of contigs ranging from 8 to 11, except for *C. metrosideri* (CBS 16091) with an assembly of 17 contigs (Table 1). Like *C. auris*, most assemblies include 7 large contigs and an 8^th^ circular contig. This observation indicates species of the *C. haemulonii* complex share the same number of chromosomes with *C. auris*. Comparison of genome sizes and gene counts revealed all species have very similar genome sizes ranging between 12.13Mb in *C. auris* CBS 12373 and 13.61Mb in *C. haemulonii* var. *vulnera* CBS 12437. *C. auris* strains tend to have slightly smaller genomes with an average of 12.4 ± 0.18Mb compared to 13.1 ± 0.34Mb for species of the *C. haemulonii* complex (Table 1).

Strains of *C. auris* are known to be highly identical, especially within clades more than >99% average pairwise nucleotide identity (ANI) was observed. But even between clades an average identity of 98.7% was reported (Muñoz et al., 2018). Here we observed similar percentages, except for *C. auris* Clade V which shared 97% identity (Supplementary Data 1). Compared to its relatives of the *C. haemulonii* complex *C. auris* shared 74% identity on average. Within the *C. haemulonii* complex, species display high similar intraspecies nucleotide identities of >99%, but on average between species 80% identity was observed (Supplementary Data 1). On the other hand, sibling species such as *C. pseudohaemulonii* and *C. duobushaemulonii* will have higher ANI’s, mostly >90%. In contrast, the distantly related species *C. albicans* and *N. glabratus* only shared an average pairwise nucleotide identity of 67-68% with *C. auris* and the other members of the *C. haemulonii* complex.

### Phylogeny of *C. auris* and its relatives of the *C. haemulonii* complex

*C. auris* is most closely related to the *C. haemulonii* species complex. After reclassification the *C. haemulonii* species complex was described to consist of *C. haemulonii*, *C. duobushaemulonii* and *C. pseudohaemulonii* and the variety *C. haemulonii* var. *vulnera* (Cendeja-Bueno et al., 2012). Recently the emerging pathogen *C. vulturna* was added as a member (Sipiczki & Tap, 2016). However, more species are linked to the *C. haemulonii* species complex including the recently described emerging pathogen *C. khanbhai* (de Jong et al., 2023) and non-pathogenic species like *C. heveicola* (Wang et al., 2008), *C. konsanensis* (Sarawan et al., 2013), *C. chanthaburiensis* (Limtong et al, 2010), *C. ruelliae* (Saluja et al., 2008), *C. ohialehuae*, and *C. metrosideri* (Klaps et al., 2021) (Table 1). Previous genomic studies only focused on the emerging human pathogens of the *C. haemulonii* complex (Muñoz et al., 2018; Gade et al., 2020). Therefore, available genomic data on the non-pathogenic species is scarce and does not provide a complete representation of the *C. haemulonii* complex. To resolve the phylogenetic relationships of species within the *C. haemulonii* complex the generated complete genomes were used to estimate the phylogeny relative to *C. auris* and other species of the *Saccharomycetales* including *C. albicans* and *N. glabratus*.

OrthoFinder was used to build a phylogenetic tree by Species Tree inference from All Genes (STAG). This method was shown to have higher accuracy than other commonly used methods, including concatenated alignments of protein sequences (Emms & Kelly, 2019). The resulting species tree placed *C. khanbhai* as well as the non-pathogenic species within the *C. haemulonii* complex (Figure 1). Interestingly, within the newly resolved phylogeny of the *C. haemulonii* complex, several ‘sub-clades’ can be observed. *C. auris* strains form a sub-clade (Figure 1) that seems to be more recently diverged based on the short branch lengths as previously reported (Muñoz et al., 2018). *C. ruelliae* appears as a more basally branching species. Next, *C. haemulonii* and *C. haemulonii* var. *vulnera* strains form a well-supported sub-clade that we call the ‘*haemulonii* clade’. Interestingly all non-pathogenic species, except one, fall within a separate sub-clade named the ‘*haemulonii* I clade’. Only *C. metrosideri* groups in the ‘*haemulonii* II clade’ which includes the known emerging pathogens *C. khanbhai*, *C. vulturna*, *C. pseudohaemulonii* and *C. duobushaemulonii* (Figure 1).

**Figure 1.**
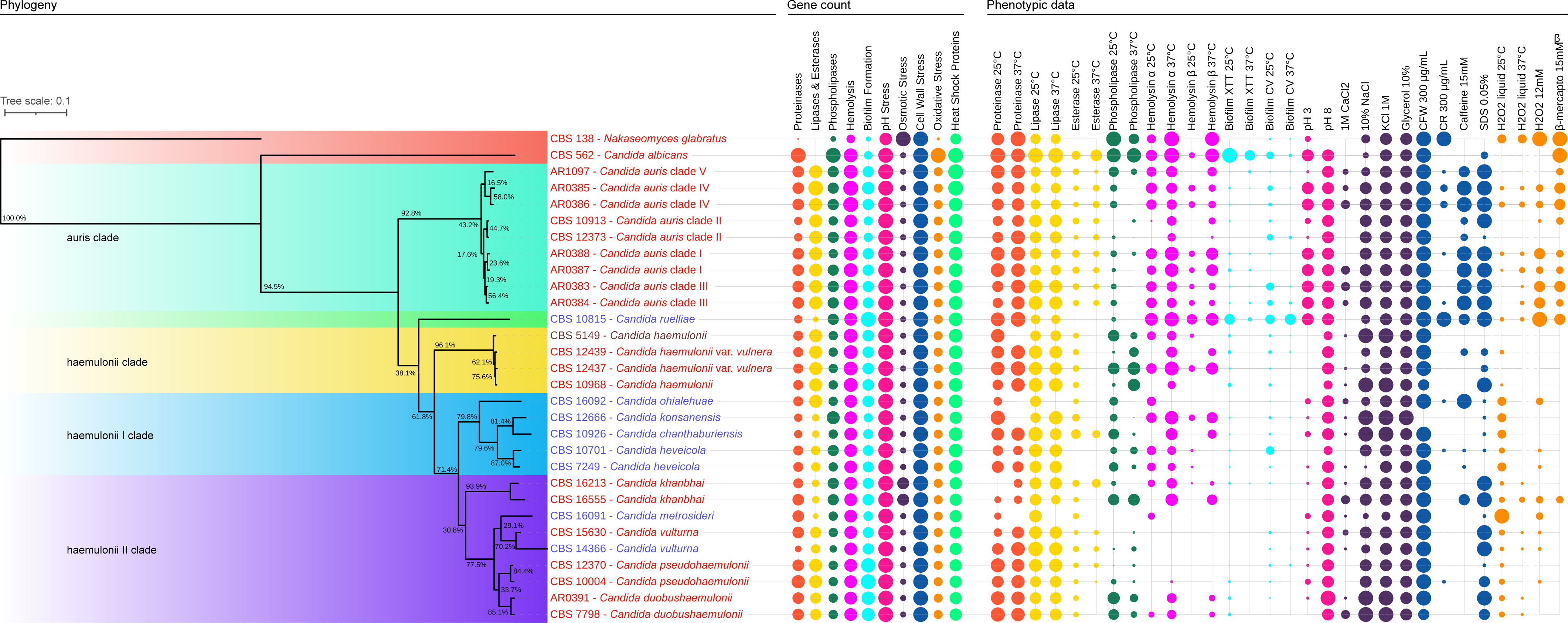
Annotated phylogeny of *C. auris* and the *C. haemulonii* complex. Phylogenetic species tree inferred by OrthoFinder using tree inference from all genes (STAG) of 30 annotated genomes including *C. auris* Clades I-V, *C. haemulonii* complex species and the distantly related species *C. albicans* and *N. glabratus*. support values at each node represent the proportion of gene-trees in which the same bipartition is found. Branch lengths indicate the mean number of changes per site. The background is color coded per identified sub-clade. Species names are color-coded, red refers to a clinical origin, brown to a veterinary origin, and blue to an environmental origin. Gene count of virulence related gene families is grouped per virulence trait and visualized next to the phylogenetic tree together with the related phenotypic data using ITOL (Letunic et al., 2021). The size of each dot represents gene family expansions or contractions and increased or decreased expression of the related phenotypes. Dots are color coded to match Gene families to their respective phenotypic test. Gene names and orthologues in *C. auris* used for the gene content analysis of each virulence trait can be found in Supplementary Data 2. Absolute values and more details about each phenotypic test can be found in the Materials and Methods and Supplementary Data 4.

### Conservation of virulence genes within the *C. haemulonii* complex

We used Helixer for gene structural annotation of the assembled genomes. Helixer improves predictions by combining traditional *de novo* hidden Markov models with deep learning (Holst et al., 2023). This resulted in nearly complete genomes. BUSCO analysis of the predicted gene sets by analysing the representation of core eukaryotic genes showed that the genomes were highly complete with on average 95.7% of these genes present in all assemblies (Table 1; Simão et al., 2015).

In line with the genome sizes, the predicted numbers of protein-coding genes were very similar among *C. auris* and the related species of the *C. haemulonii* complex, ranging from 5,629 genes in *C. auris* CBS 12373 to 6,002 in *C. haemulonii* var. *vulnera* CBS 12347. *C. vulturna* CBS 14366 showed an increased gene number of 6,174 genes, probably due to a high number of fragmented genes, shown by the BUSCO analyses (Table 1). *C. albicans* did have a similar gene number of 6,183 genes, were *N. glabratus* CBS 138 had a much lower gene count of 4,560 genes, caused by an apparent incomplete genome assembly regarding the only 85% complete BUSCOs (Table 1).

Examining the orthologous genes in *C. auris* and the *C. haemulonii* complex strains showed that the *C. auris* clades had 51 unique orthogroups compared to the close relatives of the *C. haemulonii* complex, and the more distantly related pathogens *C. albicans* and *N. glabratus* (Supplementary Data 1). Functional analyses of these orthogroups using the Interpro database (Paysan-Lafosse et al., 2023), showed most genes within these orthogroups did not contain any relevant functional protein family domains that could predict a role in the virulence of *C. auris* (Supplementary Data 1). However, three out of 51 unique orthogroups were found to contain genes that had predicted metallopeptidase, deaminase, and aspartic type endopeptidase activity. Additional PFAM analysis by HMMscan (Potter et al., 2018), predicted the presence of genes with functional domains for xylanase activity, more related to phytopathogenesis, and ferrous iron transporter proteins. Other domains detected were involved in regular cellular processes such as actin remodelling and ribosomal activity or linked to uncharacterized proteins (Supplementary Data 1)

We also examined whether gene families associated with *Candida* virulence were conserved in the *C. haemulonii* complex. *C. auris* orthologs of gene families predicted to be involved in several virulence traits such as lytic enzyme production (proteinases, lipases, and phospholipases) (Muñoz et al., 2018), hemolysis (Pendrak et al., 2004), biofilm formation (Kean et al., 2018) and stress resistance (pH, osmotic, cell wall and heat stress) (Day et al., 2018; Kim et al., 2019; Ismail et al., 2022) were identified across all species (Figure 1, Supplementary Data 2). Generally similar numbers of virulence genes are present in *C. auris* and members of the *C. haemulonii* complex. Notably, higher numbers of genes encoding the important lytic enzyme family of lipases (LIPs), were more often observed in the pathogenic species compared to the non-pathogenic species (Figure 1, Supplementary Data 2). On the other hand, similar numbers of phospholipases were present in all species. An interesting exception was found for strains of *C. auris* clade II (CBS 10913 and CBS 12373) which had a reduced number of secreted aspartyl proteases (SAPs) and also less genes involved in hemolysis and biofilm formation compared to the other *C. auris* strains.

Increased or decreased numbers for genes involved in biofilm formation could be attributed to specific gene families. Where most genes were present in similar numbers, an increased number of ALS cell surface proteins, involved in cell adhesion, was found in strains of *C. pseudohaemulonii* (7-8 genes) and *C. ruelliae* (7 genes). In contrast, in *C. auris* only 3-4 ALS genes were present and specifically in Clade II strains only 2 ALS genes were found. Moreover, numbers of *HYR*/*IFF* genes, which encode for another important adhesin-like cell surface protein family, also varied a lot. The highest numbers were found in *C. pseudohaemulonii* (14-15 genes) and *C. ruelliae* (16 genes), which was similar to *C. albicans* (15 genes). *C. auris* strains contained 8-10 *HYR*/*IFF* genes with again a decreased number in *C. auris* Clade II strains (3 genes) (Figure 1, Supplementary Data 2).

### Drug resistance within *C. auris* and the *C. haemulonii* complex

*C. auris* and its pathogenic relatives of the *C. haemulonii* complex are notorious for their multi-drug resistance, especially against azoles and to a lesser extent the echinocandins and amphotericin B (Cendejas-Bueno et al., 2012). Antifungal susceptibility testing showed this pattern of drug resistance is conserved among all species of the *C. haemulonii* complex (Table 2). All strains in this study displayed similar patterns of drug resistance. Increased resistance for fluconazole, amphotericin B and most of the triazoles (voriconazole, posaconazole, itraconazole) was common. Resistance against the echinocandins tested was rarely observed. Only after 48h of incubation *C. auris* Clade III and V strains do have increased MIC values for micafungin. For anidulafungin high MIC values are more often observed not only among *C. auris* but also within all sub-clades of the *C. haemulonii* complex.

**Table 2.** Antifungal susceptibility profiles of Candida auris and the Candida haemulonii complex. a: Minimum inhibitory concentration (MIC) to Amphotericin B (AMB), fluconazole (FLU), itraconazole (ITR), posaconazole (POS), voriconazole (VOR), isavuconazole (ISA), anidulafungin (AND), and micafungin (MCF); b: Growth only after 48 hours - as described by Cendejas-Bueno et al. (2012). Rose highlighted cells indicate MIC values above the tentative breakpoints for *C. auris* (CDC, 2018).

| Sub-clade | Strain | MIC (µg/mL)<sup>a</sup> |  |  |  |  |  |  |  |  |  |  |  |  |  |  |  |
| --- | --- | --- | --- | --- | --- | --- | --- | --- | --- | --- | --- | --- | --- | --- | --- | --- | --- |
|  |  | AMB<sup>d</sup> |  | FLU<sup>d</sup> |  | ITR |  | POS |  | VOR |  | ISA |  | AND<sup>d</sup> |  | MCF<sup>d</sup> |  |
|  |  | 24h | 48h | 24h | 48h | 24h | 48h | 24h | 48h | 24h | 48h | 24h | 48h | 24h | 48h | 24h | 48h |
| ruelliae | CBS 10815 | 1 | 2 | 16 | 32 | 0.063 | 0.25 | 0.016 | 0.063 | 0.016 | 0.031 | 0.016 | 0.063 | 0.5 | 4 | 0.125 | 0.25 |
| haemulonii | CBS 10968 | 2 | 2 | 8 | >64 | 4 | >4 | 4 | >4 | >4 | >4 | 2 | >4 | 0.063 | 0.063 | 0.063 | 0.063 |
|  | CBS 5149<sup>c</sup> | N/A | 1 | N/A | 16 | N/A | 4 | N/A | 0.125 | N/A | 4 | N/A | >4 | N/A | 0.031 | N/A | 0.063 |
|  | CBS 12437<sup>c</sup> | N/A | 1 | N/A | >64 | N/A | 0.5 | N/A | 2 | N/A | >4 | N/A | >4 | N/A | 0.063 | N/A | 0.063 |
|  | CBS 12439<sup>c</sup> | N/A | 1 | N/A | >64 | N/A | 0.5 | N/A | 4 | N/A | >4 | N/A | >4 | N/A | >4 | N/A | 0.125 |
| haemulonii I | CBS 16092<sup>b</sup> | N/A | >8 | N/A | 4 | N/A | 0.06 | N/A | 0.03 | N/A | 0.06 | N/A | N/A | N/A | 1 | N/A | 0.5 |
|  | CBS 12666 | 1 | 2 | 32 | >64 | 0.031 | 0.063 | <0.125 | 0.031 | 0.031 | 0.063 | 0.031 | 0.031 | 0.125 | 0.125 | 0.031 | 0.031 |
|  | CBS 10926 | 1 | 2 | 32 | >64 | 0.063 | 0.063 | 2 | >4 | 0.031 | 0.063 | 0.5 | 4 | 0.125 | 0.125 | 0.031 | 0.031 |
|  | CBS 7249 | 1 | 2 | 32 | >64 | >4 | >4 | 0.25 | >4 | >4 | >4 | 0.125 | >4 | 0.063 | 0.063 | 0.063 | 0.063 |
|  | CBS 10701 | 2 | 2 | 32 | 64 | 0.125 | 0.125 | 0.031 | 0.031 | 0.031 | 0.063 | 0.063 | 0.0125 | 0.125 | 0.125 | 0.031 | 0.063 |
| haemulonii II | CBS 16555 | 2 | 2 | 32 | >64 | 0.25 | >4 | 0.125 | >4 | 0.125 | >4 | 0.063 | >4 | 0.125 | 4 | 0.063 | 0.125 |
|  | CBS 16213 | 2 | 2 | 32 | >64 | >4 | >4 | >4 | >4 | >4 | >4 | >4 | >4 | 0.25 | 4 | 0.063 | 0.125 |
|  | CBS 10004 | 1 | 2 | 16 | 16 | 0.031 | 0.063 | <0.008 | 0.016 | 0.031 | 0.063 | 0.016 | 0.125 | 4 | 4 | 0.25 | 0.25 |
|  | CBS 12370 | 1 | 2 | 32 | >64 | 0.031 | 0.063 | 0.031 | 0.031 | 0.031 | 0.063 | 0.125 | 0.125 | 0.125 | 0.25 | 0.063 | 0.063 |
|  | CBS 16091<sup>b</sup> | N/A | 4 | N/A | 2 | N/A | 0.06 | N/A | 0.03 | N/A | 0.03 | N/A | N/A | N/A | 0.25 | N/A | 0.06 |
|  | CBS 14366 | 0.25 | 0.5 | >64 | >64 | 4 | >4 | <0.125 | >4 | 0.031 | >4 | 0.016 | >4 | 0.125 | 0.125 | 0.063 | 0.063 |
|  | CBS15630 | 1 | 2 | >64 | >64 | >4 | >4 | >4 | >4 | >4 | >4 | >4 | >4 | 0.125 | 0.25 | 0.063 | 0.06 |
|  | CBS16530 | 1 | 2 | >32 | >64 | >4 | >4 | >4 | >4 | >4 | >4 | >4 | >4 | 2 | >4 | 0.063 | 0.125 |
|  | CBS 7798 | 4 | 4 | >64 | >64 | >4 | >4 | 0.5 | >4 | >4 | >4 | 4 | >4 | 1 | 2 | 0.125 | 0.125 |
| auris | AR0383 | 0.25 | 1 | >64 | >64 | >4 | >4 | >4 | >4 | >4 | >4 | >4 | >4 | 0.125 | >4 | 0.5 | >4 |
|  | AR0384 | 0.5 | 1 | >64 | >64 | >4 | >4 | >4 | >4 | >4 | >4 | >4 | >4 | 0.125 | >4 | 0.5 | >4 |
|  | AR0385 | 1 | 1 | >32 | >64 | 0.5 | 0.5 | 0.25 | 0.25 | >4 | >4 | 4 | 4 | 2 | 2 | 0.5 | 1 |
|  | AR0386 | 1 | 1 | >32 | >64 | 0.5 | 0.5 | 0.125 | 0.125 | >4 | >4 | 4 | 4 | 1 | 2 | 0.25 | 1 |
|  | AR0387 | 1 | 1 | 32 | >64 | 0.063 | >4 | 0.016 | 4 | 0.063 | >4 | 0.016 | >4 | 1 | 2 | 0.125 | 0.25 |
|  | AR0388 | 1 | 2 | >32 | >64 | 0.5 | 0.5 | 0.25 | 0.25 | >4 | >4 | 2 | 2 | 2 | >4 | 0.25 | 1 |
|  | CBS 10913 | 0.5 | 1 | 16 | 32 | 0.031 | 0.063 | <0.008 | 0.016 | 0.016 | 0.031 | <0.008 | 0.016 | 0.031 | 0.031 | 0.031 | 0.031 |
|  | CBS 12373 | 0.5 | 1 | >32 | >64 | 0.25 | 0.25 | 0.125 | 0.25 | 0.5 | 4 | 1 | 2 | 0.063 | 1 | 0.063 | 0.125 |
|  | AR1097 | 1 | 1 | >32 | >64 | 0.5 | 0.5 | 0.25 | 0.25 | 1 | 4 | 1 | 4 | 0.125 | 1 | 0.125 | >4 |

<sup>b</sup> Growth only after 48 hours - as described by Cendejas-Bueno et al. (2012).
<sup>c</sup> CBS 16091 and CBS 16092 maximum growth temperature is 30°C - therefore data retrieved from original article by Klaps et al. (2020).
<sup>d</sup> Pink highlighted cells indicate strains with MIC values above tentative breakpoints for *C. auris* (CDC, Atlanta, GA, USA): AMB ≥2 µg/mL, FLU ≥32 µg/mL, AND ≥4 µg/mL, and MCF ≥4 µg/mL.

Due to the worrying ability of *C. auris* to develop resistance against all available classes of antifungal drugs, the mechanisms associated with drug resistance have been well studied in *C. auris* (Rybak et al., 2022). We identified orthologs of genes noted to confer drug resistance in *C. auris*, in all species of the *C. haemulonii* complex (Supplementary Data 3). Multiple sequence alignment showed that most of these genes are well conserved within the *C. haemulonii* complex. Interestingly, none of the drug-resistant mutations described specifically for *C. auris* (Rybak et al., 2022) were observed in the other species (Supplementary Data 3). Only in the important azole target *ERG11* several commonly described mutations were found, also reported to cause drug resistance in *C. albicans* for example (Lockhart et al., 2017). This included mutations F105L, S110A, D116A, D153E, R267T, and A432S, which were detected in all the sub-clades and almost all strains (Supplementary Data 3) (Lockhart et al., 2017). Other important drug targets like the echinocandin target *FKS1* or the amphotericin B target *ERG6* were well conserved, but did not contain any of the described mutations, indicating other mechanisms are causing the observed resistance against these classes of drugs in the *C. haemulonii* complex.

Therefore, orthologs of transporters involved in clinical antifungal resistance in *C. albicans* and *C. auris* were also identified (Supplementary Data 2). Previously high copy numbers of oligopeptide transporters (OPT) and siderophore iron transporters (SIT) were reported in *C. auris* and the *C. haemulonii* complex (Muñoz et al., 2018). We observed similar numbers for these transporters in all species of the *C. haemulonii* complex. Strains of the *C. haemulonii* sub-clade had the highest number with 14 and 4 copies of OPT and SIT genes, respectively. Additionally, transporters from the ATP binding cassette (ABC) and major facilitator superfamily (MFS) may confer resistance by overexpression (Muñoz et al., 2018). We identified orthologs of the ABC transporter family CDR (*CDR1*, *CDR2*, *CDR4*, and *CDR11*) and *SNQ2* across the whole *C. haemulonii* complex. Three genes related to the CDRs were found in all strains. For *SNQ2* all *C. auris* strains contained two related genes, whereas most of the *C. haemulonii* complex species had 1 or 2 related genes. Strains of the non-pathogenic species *C. heveicola* (CBS 10701 and CBS 7249) as well as *C. konsanensis* (CBS 12666) and *C. chanthaburiensis* (CBS 10926) were lacking a *SNQ2* ortholog. On the other hand, only one copy of the MFS transporter *MDR1* was found in *C. auris*, where all other species contained two copies. *C. ruelliae* (CBS 10815) and *C. vulturna* (CBS 14366) even contained three *MDR1* copies. Finally, orthologs of the transcription factors *TAC1B*, and *MRR1* which regulate the expression of the ABC and MFS transporter families were also detected (Rybak et al., 2020; Li et al., 2022). Nevertheless, multiple mutations in *TAC1B* that lead to overexpression of the CDR genes were only found in the *C. auris* strains. Also, the N647T mutation in *MRR1*, described to cause overexpression of *MDR1*, was only observed in *C. auris* Clade III strains (Li et al., 2022).

### Phenotypic differences related to virulence between *C. auris* and the *C. haemulonii* complex

We observed limited genetic diversity between *C. auris* strains and those of the *C. haemulonii* complex. Therefore, we explored whether specific phenotypic variation could be observed among the strains, specifically focussing on infection relevant traits. Physiological characterization did not show variation that could be linked to a more virulent phenotype (Supplementary Data 4). Also, between the sub-clades a specific physiological profile was not observed. Some intra- and inter-species variability was found regarding growth on different carbon and nitrogen sources but did not vary between pathogenic and non-pathogenic strains. However, compared to the distantly related pathogens *C. albicans* and *N. glabratus*, a greater diversity regarding assimilation and fermentation of different carbon sources was observed within *C. auris* and the *C. haemulonii* complex. The most significant difference was the ability to grow at high temperatures. Only *C. auris*, *C. ruelliae*, *C. khanbhai* and *C. albicans* were able to grow at 42°C. However, all species except for *C. metrosideri* and *C. ohialehuae* were able to grow at human body temperature of 37°C.

Lytic enzymes are an important virulence factor allowing the fungus to invade and survive within the host (de Jong & Hagen, 2019). Despite the increased number of genes encoding lipases in the genome of *C. auris* and other emerging pathogens of the *C. haemulonii* complex, no increased activity of these enzymes was found compared to the non-pathogenic species (Figure 1). Esterase activity was more present in the *C. auris* strains at 37°C, but high activity was also found for the non-pathogenic species *C. chanthaburiensis* and *C. heveicola*. Hemolytic activity was more common in *C. auris* compared to most species. Especially increased β-hemolysis was observed within the *C. auris* strains of Clade I at 37°C. On the other hand, some non-pathogenic species such as, *C. ruelliae*, *C. konsanensis* and *C. chanthaburiensis* displayed similar hemolytic activity. In general, the common pathogens *C. albicans* and *N. glabratus* had the highest lytic enzyme activities. Especially, regarding phospholipase and β-hemolytic activity.

Biofilm formation is seen as another important virulence factor of *Candida* species. Biofilm formation is linked to increased antifungal resistance and protects the cells against to host immune system (de Jong & Hagen, 2019). Yet, low levels of biofilm formation were observed in almost all *C. auris* and *C. haemulonii* complex strains. Compared to *C. albicans*, only *C. ruelliae* produced similar biofilms, with more biomass (crystal violet staining) but less metabolic activity (XTT reduction). *C. ruelliae* did have an increased number of genes involved in biofilm formation, but so did strains of *C. pseudohaemulonii*, which lacked increased biofilm formation under the conditions tested.

Ultimately, the most significant phenotypic differences were observed regarding traits involved in stress resistance. The ability to adapt to stresses imposed by the host during invasion is required for the pathogenicity of most fungal pathogens (Day et al., 2018; Heaney et al., 2020). Where all strains could grow at pH 8, only *C. auris, C. ruelliae* and *C. albicans* would have good growth at the more acidic pH 3. High resistance to cationic stress imposed by either sodium chloride (NaCl) or calcium chloride (CaCl_2_) was observed among all *C. auris* and *C. haemulonii* complex strains. Some strains even displayed improved growth in the presence of NaCl, including strains of *C. haemulonii*, *C. duobushaemulonii*, C. *konsanensis* and *C. chanthaburiensis*. Growth on CaCl_2_ was more restricted, but most *C. auris* and *C. haemulonii* complex strains did show some growth on plates containing CaCl_2_, where *N. glabratus* and *C. albicans* did not grow. *C. auris* also seems best adapted to cell wall stress (calcofluor white, Congo red, caffein, SDS). Especially on caffein medium *C. auris* strains displayed better growth compared to most species. Only *C. ruelliae* had better resistance, growing in the presence of all cell wall-damaging agents. Inducing oxidative stress by exposure to hydrogen peroxide (H_2_O_2_) showed an interesting pattern. Most non-pathogenic species were able to grow at high H_2_O_2_ concentrations, but only at 25°C.

Increasing the temperature to 37°C, strongly reduced the H_2_O_2_ concentration at which these species could grow. Only *C. auris*, *C. vulturna*, *C. khanbhai*, *C. ruelliae* and *N. glabratus* grew at similar or even higher concentrations at 37°C. Notably, strains of *C. auris* clade II and V and *C. albicans* would not grow under any oxidative stress. In addition, the response to endoplasmic reticulum (ER) stress was observed by growth on β-mercapto-ethanol. *C. albicans* and *N. glabratus* grew well under ER stress, where most *C. haemulonii* complex strains did not show any growth. The exceptions were *C. auris*, *C. ruelliae* and *C. khanbhai* (CBS 16555).

### *In vivo* virulence of *C. auris* and related species of the C. *haemulonii* complex

To evaluate the pathogenicity of *C. auris* compared to its relatives of the *C. haemulonii* complex we used the invertebrate *Galleria mellonella* fungal infection model. All strains were able to cause death, but mostly at low levels. *C. auris* strains showed to be significantly more virulent (Figure 2; Supplementary Data 5). However, *C. auris* strains displayed high inter clade variability. Strains belonging to Clade I were most virulent compared to the other *C. auris* strains and were even more virulent than *C. albicans*. Strains of Clade IV displayed similar virulence as those of Clade I, where strains of Clade III and especially Clade II were significantly less virulent (Figure 2, Supplementary Data 5). Of the *C. haemulonii* complex only *C. ruelliae* showed increased virulence, killing similar amounts of larvae as *C. auris* clade II and III strains. Interestingly, the other known emerging pathogens *C. haemulonii*, *C. vulturna*, *C. pseudohaemulonii*, *C. khanbhai* and *C. duobushaemulonii*, did not have any increased virulence compared to their non-pathogenic relatives.

**Figure 2.**
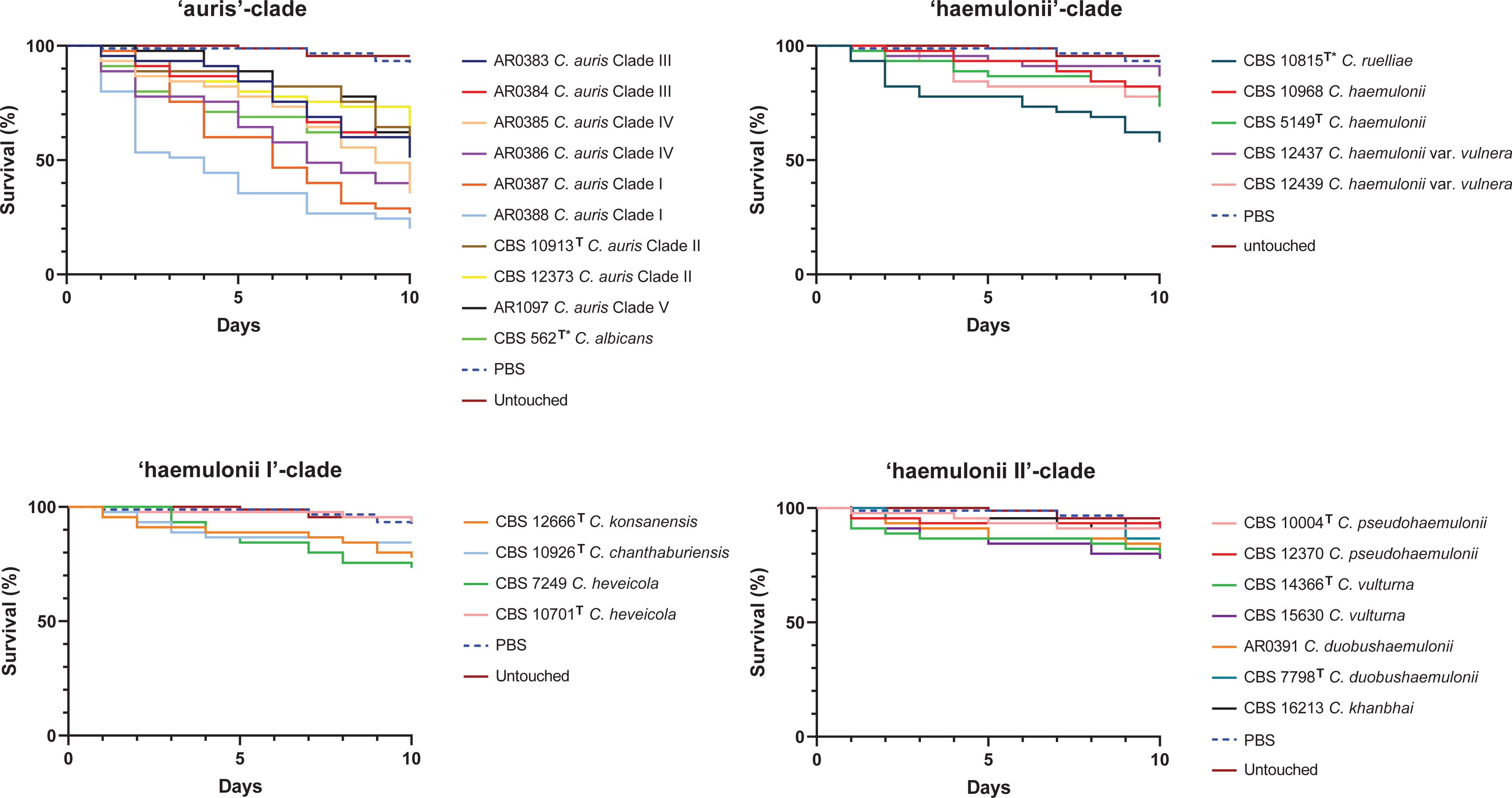
The virulence of *C. auris* and its relatives of the *C. haemulonii* complex in *Galleria mellonella* larvae at 37 °C. Kaplan–Meier survival plots of *Galleria mellonella* injection with 10^6^ CFU/larva of the indicated *Candida* species, organized by their sub-clade; *C. auris* (A), ‘haemulonii’ (B), ‘haemulonii I’ (C), and the ‘haemulonii II’ (D) are shown. Equivalent plots obtained with phosphate-buffered saline injected larvae (PBS) and untouched larvae are included in all four panels as controls. Per strain 15 larvae were inoculated. Experiments were performed in triplicate; plots represent the combined (additive) data from all experiments (n = 45). Statistical comparison of each survival curve (pairwise Log-rank test: P<0.05; ns, P>0.05) can be found in Supplementary Data 5. \**C. albicans* (CBS 562) was added as a reference strain to the *C. auris*-clade plot and *C. ruelliae* (CBS 10815) was added to the haemulonii-clade plot for comparison.

## Discussion

The rise of *Candida auris* showed the world the ability of fungi to rapidly become an urgent threat to public health (Lockhart et al., 2017). The worsening spread of *C. auris* and the simultaneous emergence of its close relatives of the *C. haemulonii* complex (Geddes-McAlister et al., 2019; Garcia-Solache et al., 2010; Denning et al., 2024), require a thorough understanding of the virulence traits of these pathogens in order to stop their spread and initiate optimal treatment. Comparing pathogens to their closest non-pathogenic relatives is a common strategy to provide new insights into how virulence appears and evolves across species. Proven in bacteria and fungi (Moran et al., 2011; Wurtzel et al., 2012), including *Candida* (Butler et al., 2009), this strategy is promising to generate valuable insights into the virulence of *C. auris* and the *C. haemulonii* complex. Here, we generated new genomic and phenotypic data for all *C. auris* clades, and both pathogenic and non-pathogenic relatives within the *C. haemulonii* complex. We observed that *C. auris* and its relatives of the *C. haemulonii* complex are genetically very similar and harbour traits that likely prime them to become human pathogens. Despite the shared genetic virulence properties, *C. auris* was shown to be significantly more virulent in the *G. mellonella* fungal infection model. Our phenotypic analysis highlights unique traits of *C. auris* compared to its *C. haemulonii* complex relatives that likely contributed to this increased virulence. Overall, this work lays a foundation to study the evolution of virulence in *C. auris* and its close relatives.

Genomic comparison of *C. auris* to its siblings was impaired by the lack of genomes available for most species of the *C. haemulonii* complex. Using nanopore long-read sequencing we generated highly continuous and complete genomes for all described relatives of *C. auris,* including the non-pathogenic ones. With these annotated genomes the full current phylogenetic relationship of the *C. haemulonii* complex could be resolved. Our phylogenetic analysis showed that the *C. haemulonii* complex consists of several sub-clades. In line with previous studies, *C. haemulonii* is more distantly related to the other emerging species *C. vulturna*, *C. pseudohaemulonii* and *C. duobushaemulonii*, which here form a sub-clade with the newly described pathogen *C. khanbhai* (Muñoz et al., 2018; Gade et al., 2020; Francisco et al., 2023). Interestingly, all non-pathogenic species group together, except for *C. metrosideri* –which groups with *C. vulturna*– and *C. ruelliae* that is the basal branching species. The resolved phylogeny highlights the repeated evolution of virulence within the *C. haemulonii* complex.

Repeated evolution of virulence suggests that *C. auris* and its relatives harbour traits that pre-adapt them to become human pathogens. *C. auris* and its pathogenic relatives *C. haemulonii*, *C. duobushaemulonii*, and *C. pseudohaemulonii*, share notable expansions of virulence related gene families described in *C. albicans* and other common pathogenic *Candida* species (Muñoz et al., 2018; Butler et al., 2009). Notably, the shared expansion of lipases between all pathogens suggests a shared mechanism of virulence (Muñoz et al., 2018). Here we observed that this expansion of lipases is indeed present in *C. auris* and most pathogens of the *C. haemulonii* complex, supporting the idea that lipases play an important role in the evolution of virulence within *Candida*. In contrast, the non-pathogenic species *C. ohialehuae* exhibited a similar number of lipases, suggesting the expansion of lipases alone is not enough to become pathogenic. This idea was supported by *C. auris* Clade II strains, that do share the same number of lipases with other pathogens, but are significantly less virulent in *in vivo* infection models (Forgács et al., 2020; Abe et al., 2020). Moreover, *C. auris* Clade II strains have a seemingly unique propensity for the ear instead of invasive infection (Welsh et al., 2019). Notably, a significant reduction of *HYR*/*IFF* adhesins from 8 to 3 copies was found in these Clade II strains, which was previously reported to be caused by the deletion of sub-telomeric regions enriched in these and other cell-surface proteins (Muñoz et al., 2021). The reduced virulence of Clade II strains was linked to this loss of cell-wall proteins (Muñoz et al., 2021). In addition, our analysis showed the loss of SAP genes encoding secreted proteases, which fulfil multiple specialized functions during *Candida* infection (Naglik et al., 2003). In contrast, *C. auris* clades did possess unique genes with proteolytic activity. On the other hand, these were also present in *C. auris* clade II strains which makes it unlikely that these gene will play a major role in the virulence of *C. auris*. Since the non-pathogenic species, shared gene content for most of the analysed virulence related gene families, they indeed seem primed to become human pathogens. However, most of the virulence gene orthologs identified in *C. auris* and the other species still need to be characterized as their function has been mainly inferred from the distantly related *C. albicans* (Muñoz et al., 2018). Moreover, we did not take genomic re-arrangements, and discrete point mutations in coding or non-coding regions into consideration that may also be key factors in the evolution of virulence.

*C. auris* and its relatives are notorious for their multi-drug resistance. Resistance mechanisms of *C. auris* have been studied thoroughly (Rybak et al., 2022), but not much is known about those of its close relatives of the *C. haemulonii* complex. The shared antifungal resistance phenotype between *C. auris* and most of the siblings in the *C. haemulonii* complex indicates that common resistance mechanisms are present. Nevertheless, resistance mutations described for *C. auris* in the well conserved azole and echinocandin drug targets, *ERG11* and *FKS1*, respectively (Rybak et al., 2022), were not found in any of the other closely related species. Also, the recently reported mutation YY98V* in *ERG6*, which could have explained the generally reduced sensitivity for amphotericin B, was not detected (Rybak et al., 2022). The lack of known drug resistance mutations in members of the *C. haemulonii* species complex pointed out that other molecular mechanisms likely play a role. *C. auris* has the unique ability to rapidly develop drug resistance either by acquiring point mutations, increasing transcription, or copy number variation (Bassetti et al., 2019; Lockhart et al., 2017). In contrast, resistance of *C. haemulonii* complex members seem to rely on more conserved intrinsic mechanisms, yet to be understood (Francisco et al., 2023). Alternatively, multiple transporter families such as oligopeptide transporters (OPT), siderophore iron transporters (SIT), ABC transporters and MFS transporters were all reported to confer resistance through upregulation during exposure to antifungals (Muñoz et al., 2018; Li et al., 2022; Wasi et al., 2019). We did see shared gene content for all these types of transporters in *C. auris* and its relatives but with variations between the sub-clades. The variability in gene content for transporter families suggests that *C. auris* and its relatives rely on different transporters during their antifungal drug response. Our genomic analyses showed the relative variation of orthologues genes within the genomes of the species. Hence, more transcriptomic data is needed to shed light on specific molecular mechanisms activated by species of the *C. haemulonii* complex during exposure to the different classes of antifungal drugs.

Our phenotypic analysis highlighted *C. auris* and its close relatives share the same physiological characteristics but do vary in some crucial virulence traits. Efficient metabolic adaptation was shown to be at the basis of *C. albicans* virulence (Brown et al., 2014). Notably, members of the *C. haemulonii* complex displayed assimilation and fermentation of even more carbon sources than *C. albicans*. Following the assimilation of local nutrients, a fungus should be able to counter any local environmental stresses and evade host defences (Brown et al., 2014; Lionakis et al., 2023). Especially when it comes to stress resistance, *C. auris* showed a unique phenotype compared to most of its relatives. This unique stress resistance phenotype probably increases the ability of *C. auris* to survive within the host and cause disease. Interestingly, *C. auris* Clade II strains were much less stress resistant. Which is in accordance with the previously mentioned propensity to only cause ear-infections (Welsh et al., 2019). It is tempting to speculate that the here observed phenotype of *C. auris* could explain its pathogenic success. However, the non-pathogenic species *C. ruelliae* had an almost identical phenotype. On the other hand, *C. khanbhai* also shared the stress resistant phenotype and has been reported as an emerging nosocomial pathogen. In the end, stress resistance is most likely a combinatorial virulence trait. Moreover, the expression of phenotypic traits *in vitro* is heavily affected by the culture conditions used. For example, *C. auris* biofilm formation was almost absent under the standard conditions in this study, where recent studies show a high biofilm forming capacity for *C. auris* when grown in synthetic sweat medium, mimicking axillary skin conditions (Horton et al., 2020; Biswas et al., 2023).

Ultimately, *in vivo* models remain essential to mimic the multifarious environment needed to study the pathogenesis of infectious diseases. *G. mellonella* larvae have been widely adopted to study fungal infections (Pereira et al., 2018). Our survival analysis showed that *C. auris* was significantly more virulent than its relatives, especially Clade I strains. This is in line with the genotypic and phenotypic observations indicating an increased virulence potential for *C. auris*. The reduced virulence of Clade II strains which lack some of these important traits also supports this direct relationship. Notably, pathogens and non-pathogens within the *C. haemulonii* complex displayed similar virulence. Substantiating the idea that these species all have a pathogenic potential. An explanation for the clinical emergence of only a sub-set of species may lay in their ecological niche. The emerging pathogens of the *C. haemulonii* complex probably reside in environments where they are easier transmitted to humans allowing them to cause infection. Despite recent sampling efforts, the ecological niches of all species including *C. auris* remain elusive (Arora et al., 2021; Escandón, 2022; Irinyi et al., 2022). Therefore, we could only speculate how these species emerge into the nosocomial environment. To this end, more environmental sampling studies should be conducted.

The data presented in this study provides a basis for further analysis of the virulence and resistance mechanisms in *C. auris* and the *C. haemulonii* complex. Unique genomic and phenotypic traits seem to have contributed to the increased virulence of *C. auris*, where shared traits have primed the whole *C. haemulonii* complex for virulence. Transcriptomic together with gene deletion studies should shed light on the role of shared and species-specific genes in virulence and resistance. As we are beginning to understand the biology of these species, future research should also focus on their ecology. More sampling is needed to find out where these species are hiding and what drives their emergence into the nosocomial environment. The data provided here could aid in the development of specific and rapid diagnostic tools to monitor the spread of this important emerging group of fungal pathogens.

## Acknowledgements

*Candida auris* reference strains were provided by the Centers for Disease Control and Prevention (Atlanta, GA, USA) Antimicrobial Resistance Isolate Bank. Authors thank the depositors of the strains that were made available by the CBS Culture Collection hosted at the Westerdijk Fungal Biodiversity Institute (Utrecht, The Netherlands).

## Author’s contributions

Auke W. de Jong: Conceptualization; Data curation; Formal analysis; Investigation; Methodology; Validation; Visualization; Roles/Writing - original draft; and Writing - review & editing. Sander Boden: Data curation; Formal analysis; Investigation; Methodology; Software; Validation; Visualization; Roles/Writing - original draft; and Writing - review & editing. Annemarie Zandijk: Formal analysis; Investigation; Methodology; Writing - review & editing. Alexandra M. Kortsinoglou: Investigation; Writing – review & editing. Bert Gerrits van den Ende: Formal analysis; Investigation; Methodology; Writing - review & editing. Elaine C. Francisco: Investigation; Methodology; Resources; Writing - review & editing. Podimata Konstantina Nefeli: Investigation; Writing – review & editing. Vassili N. Kouvelis: Investigation; Resources; Supervision; Writing – review & editing. Miaomiao Zhou: Conceptualization; Investigation; Methodology; Project administration; Resources; Supervision; Validation; Writing - review & editing. Ferry Hagen: Conceptualization; Data curation; Formal analysis; Funding acquisition; Investigation; Methodology; Project administration; Resources; Software; Supervision; Validation; Visualization; Roles/Writing - original draft; and Writing - review & editing.

---

**Supplementary Data 1**. Genome assembly statistics and orthogroup analysis of *C. auris* and the *C. haemulonii* species complex.

**Supplementary Data 2.** Gene copy number variation and conservation analysis of genes associated with drug resistance and virulence in *C. auris* and the *C. haemulonii* species complex.

**Supplementary Data 3.** Observed amino acid substitutions in *C. auris* compared with *C. auris* strains used in this study and strains of the *C. haemulonii* species complex after multiple alignment of the drug resistance genes *ERG11*, *FKS1*, *TAC1B*, *UPC2*, *FUR1*, *CIS2*, *MEC3*, *MRR1*, *ERG3*, *ERG6*, and *FUR1*.

**Supplementary Data 4.** Phenotypic characterization of *C. auris* Clade I-V strains and close relatives of the *C. haemulonii* complex, together with reference strains *C. albicans* CBS 562 and *N. glabratus* CBS 138 for comparison.

**Supplementary Data 5.** Statistical analyses of species-specific differences in pathogenicity in the *G. mellonella* fungal infection model.

